# Identification of potential selective autophagy receptors from protein-content profiling of autophagosomes

**DOI:** 10.1101/2023.03.14.532537

**Authors:** Alberto Cristiani, Arghya Dutta, Sergio Alejandro Poveda-Cuevas, Andreas Kern, Ramachandra M. Bhaskara

**Affiliations:** Institute of Biochemistry II, School of Medicine, Goethe University Frankfurt, Theodor-Stern-Kai 7, 60590 Frankfurt (Main), Germany; Buchmann Institute for Molecular Life Sciences, Goethe University Frankfurt, Max-von-Laue Str. 15, 60438 Frankfurt (Main), Germany; Institute of Pathobiochemistry, University Medical Center of the Johannes Gutenberg University, Duesbergweg 6, 55099 Mainz, Germany

**Keywords:** Computational Biology, Cell Biology, Data Analysis, Autophagosomes, Proteomics, Receptors, Molecular Models

## Abstract

Selective autophagy receptors (SARs) are central to cellular homeostatic and organellar recycling pathways. Over the last two decades, more than 30 SARs have been discovered and validated using a variety of experimental approaches ranging from cell biology to biochemistry, including high-throughput imaging and screening methods. Yet, the extent of selective autophagy pathways operating under various cellular contexts e.g., under basal and starvation conditions, remains unresolved. Currently, our knowledge of all known SARs and their associated cargo components is fragmentary and limited by experimental data with varying degrees of resolution. Here, we use classical predictive and modeling approaches to integrate high-quality autophagosome content profiling data with disparate datasets. We identify a global set of potential SARs and their associated cargo components active under basal autophagy, starvation-induced, and proteasome-inhibition conditions. We provide a detailed account of cellular components, biochemical pathways, and molecular processes that are degraded via autophagy. Our analysis yields a catalog of new potential SARs that satisfy the characteristics of bonafide, well-characterized SARs. We categorize them by the subcellular compartments they emerge from and classify them based on their likely mode of action. Our structural modeling validates a large subset of predicted interactions with the human ATG8 family of proteins and shows characteristic, conserved LC3-interacting region (LIR)–LIR-docking site (LDS) and Ubiquitin-interacting motif (UIM)–UIM-docking site (UDS) binding modes. Our analysis also revealed the most abundant cargo molecules targeted by these new SARs. Our findings expand the repertoire of SARs and provide unprecedented details into the global autophagic state of HeLa cells. Taken together, our findings provide motivation for the design of new experiments, testing the role of these novel factors in selective autophagy.

## I. INTRODUCTION

Autophagy is one of the most conserved and prevalent homeostatic pathways in eukaryotic cells. This catabolic process is under strict regulation to identify and recycle unwanted or dysfunctional cellular components in response to varying cellular stresses^1^. Macroautophagy, a dominant process in eukaryotes among the several variations over the theme, involves cytoplasmic materials sequestered into double-membrane bound vesicles for delivery to lysosomes^2^. Autophagic membranes in mammals are presumed to emerge from the ER as cup-shaped phagophores, which grow in size encapsulating cargo in double-membraned vesicles called autophagosomes^3^. These large vesicles (∼ 300–1000 nm in diameter) fuse with lysosomes for degradation of their contents by hydrolytic enzymes^4^. Autophagic pathways can be selective and non-selective depending on how cargo components are sequestered. Non-selective or bulk pathways involve direct uptake of cytoplasmic material and are nonspecific to substrates, whereas cargo selection and specificity are key determinants of selective pathways^1,5^.Selective autophagy relies on specialized proteins called cargo receptors or selective autophagy receptors (SARs)^5^. They can be soluble, or membrane-bound proteins that recognize cargo components and actively engage with the growing phagophore to promote autophagosome formation in close association with cargo sites^5,6^. Cargo-bound SARs bind to phagophore-anchored lipidated protein variants of the ATG8 protein family^7^. In humans 6 ATG8 family members have been identified^8^. This interaction is critical for sequestering the phagophore at the cargo site. It is mediated by a characteristic sequence motif with core sequence [W/F/Y]_0_–X_1_–X_2_–[L/I/V]_3_ that is housed within the SAR called the LC3-interacting region (LIR) or GABARAP interacting motif (GIM). The LIR/GIM segments (referred to as LIR hereafter) fit into two well-conserved surface hydrophobic pockets in all ATG8 proteins, called the LIR docking site (LDS)^5,7^. Ubiquin-interacting motifs (UIMs) on SARs can also bind to hATG8 proteins^9^.

The LIR–LDS binding mode between distinct SARs and ATG8 proteins forms the molecular basis of all known selective autophagy pathways^10^. In mammals, p62/SQSTM1 (sequestosome-1) was first demonstrated to directly bind to ATG8/LC3 to facilitate the degradation of ubiquitinated protein aggregates via autophagy^11^. Since then, more than 30 SARs have been discovered in humans alone. SARs can be broadly grouped into soluble and membrane-bound cargo receptors based on their cellular localization^5,12^ and classified as ubiquitin-dependent/-independent SARs based on the presence/absence of distinct ubiquitin-binding domains (UBDs)^9^.

Post-translational modifications (PTMs) regulate the action of SARs^13^. PTMs serve as a switching mechanism for activating SAR, enhancing cargo recruitment and selection, in response to signaling cascades, varying nutrient states, and stress conditions. For instance, p62/SQSTM1 is phosphorylated by CK2 and TBK1 in response to amino-acid sensing, oxidative stress, and DNA damage^14^. These modifications enhance its affinity to ubiquitinated cargos^15^. A detailed list of diverse PTMs employed for regulating SARs, including phosphorylation, ubiquitination, acetylation, SUMOylation, and UMFylation, is discussed in several recent reviews^12,16^.

Distinct subsets of SARs are employed to eliminate different cellular components^17^. Various studies have shown how mitochondria^18^, peroxisomes^19^, lysosomes^20^, endoplasmic reticulum^21^, and the nucleus^22^ are selectively degraded using dedicated SARs. Although distinct SARs have been identified for most organelles and compartments, the entire list of cellular components or organelles targeted by selective autophagy remains incomplete.

Several aspects of SARs and their associated cargo remain unclear. For example, the complete list of functional SARs in cells is still unknown, and their cargo specificities remain untested. As most studies focus on the identification and characterization of individual receptors and associated cargos, a clear global perspective is lacking. Furthermore, the confluence of several homeostatic pathways adds to the complexity of autophagic processes. Additionally, individual autophagosome formation events are ubiquitous, temporally, and spatially resolved in cells, but are often analyzed in bulk, averaging out signals specific to selective pathways.

Schmitt et al. ^23^ recently introduced an antibody-based approach using FACS to isolate high-quality native autophagic vesicles in large quantities^23^. By employing quantitative proteomics, they were able to profile the protein contents of human autophagic vesicles purified from HeLa cells under (i) basal conditions, (ii) EBSS-induced starvation, and, (iii) MG132-mediated proteasome inhibition, providing a wealth of new information. More specifically, upon proteasome inhibition, they showed treatment-specific changes in the autophagic vesicles’ protein contents, demonstrating the sensitivity of their powerful isolation-profiling approach. Based on these observations, we hypothesize that high-quality protein-content profiling of pure and intact autophagic vesicles is a valuable resource for gauging the global autophagic state of cells. Further, it can be mined to provide quantitative data on the various selective autophagy pathways and processes operating under any given cellular context.

Here, we revisit the proteomics datasets collected by Schmitt et al. ^23^ By integrating disparate information from multiple external repositories and combining them with new predictive analyses; we highlight the role of selective autophagy pathways in determining autophagosomal protein contents. We identify new potential SARs and their associated cargo components. We quantify the contribution of specific SARs and their plausible functions toward the total autophagic flux under different cellular conditions.

## II. METHODS

### A. Datasets

We used the proteomics dataset provided in Schmitt et al^23^ (PRIDE Project: PXD024419)^24^ to obtain the protein inventory of autophagosomes. It contains read counts of 4627 proteins from isolated autophagosomes under three conditions—basal autophagy, starvation-induced (EBSS), and proteasome inhibition (MG132)—each of which was obtained from three experimental replicates. Due to the ambiguous mapping of peptides to multiple proteins in the initial dataset, we remapped all the proteins entries into an extended dataset (PTMX) corresponding to 5273 unique UniProt accession codes. Quantile normalization was used to normalize the read counts in each replicate. The relative abundances of proteins under EBSS treatment compared to the Basal condition were calculated as log_2_(avg.(*EBSS*)*/*avg.(*Basal*)), similarly for MG132, with the averages calculated across three replicates. The significance of the over-/under-representation of proteins in a given condition was calculated by comparing condition and basal replicas using a two-sided *t*-test and represented as volcano plots. A gold standard positive (GSP) dataset of proteins with experimentally validated ATG8-interacting LIR motifs was compiled from literature^25–27^. Based on a critical analysis of these proteins, we categorized 41 proteins as GSP SAR+ with established roles in selective autophagy. The remaining GSP proteins were categorized as GSP SAR– due to untested roles in selective autophagy and few examples corresponding to the basic autophagic machinery. In total, the GSP dataset comprises 120 proteins.

### B. LIR site predictions

LIR sites in PTMX (44 847) were identified by scanning sequences using iLIR^28^. We also tested iLIR predictions for experimentally validated LIR sites corresponding to proteins in GSP datasets. We found 30 LIR sites spanning 23 SAR+ proteins and 73 LIR sites spanning 58 SAR– proteins using iLIR. We used a PSSM cut-off of 16 to identify high-confidence LIR sites within PTMX proteins and identified 1892 sites in 1401 proteins^28,29^. These 1401 proteins constitute the potential SARs (pSARs) dataset.

### C. Functional clustering of autophagosomal proteins

Functional clustering of proteins present in the PTMX dataset was performed after annotating them with GO terms corresponding to three different ontologies, followed by enrichment analysis and comparison with the human proteome using DAVID^30^ with default parameters. The top-ranked enriched clusters were represented for each ontology, along with the number of proteins and relative abundance. DAVID analyses were performed for: (i) 3000/4464 most abundant overlapping proteins; (ii) 500 most abundant proteins detected under basal, EBSS- and MG132-treatments, independently; and (iii) 1401 newly identified pSAR proteins.

### D. Mapping cellular localization of pSARs

The primary subcellular location of all 1401 pSAR proteins was compiled from the Human Protein Atlas (https://www.proteinatlas.org/) after filtering experimental evidence type “approved”, “supported,” or “enhanced”^31^. The pSARs were categorized into UBD+, and UBD– based on the presence/absence of at least one of the 20 well-known ubiquitin-binding (UBD) domains with unique InterPro accession IDs^32,33^. The pSARs were categorized into TM+, and TM– based on the presence/absence of at least one consensus trans-membrane segment as annotated in the Human Transmembrane Proteome database^34^.

### E. Mapping ATG8 interactions and PTMs of pSARs

We retrieved a dataset of pSAR–hATG8 interactions by combining the STRING^35^ and BioGRID^36^ repositories in PSI-MI format. We filtered for direct physical pairwise interactions using the PSI-MI terms MI:0915 (physical association), MI:0407 (direct interaction), and MI:0914 (association). We used the same filtering process to obtain pSAR–PTMX interactions to identify interacting cargo components. A dataset of experimentally validated PTMs (phosphorylation, ubiquitination, sumoylation, acetylation)—that are proximal to predicted LIR sites (PSSM *≥* 16) of pSARs—was mapped from data collected from PhosphoSitePlus and dbPTM^37,38^.

### F. Clustering of pSARs and SAR+

To cluster the pSAR and SAR+ proteins, we first performed pairwise global alignments using a BioPython implementation of Needleman–Wunsch algorithm (Align module)^39,40^ and computed the pairwise gamma distance given by *d*_*γ*_ = *a*[(1*− p*)^*−*1*/a*^*−* 1], where *a* = 2 and *p* = *n*_d_*/n* in which *n*_d_ is the number of mismatches and *n* is the alignment length^41^. Protein clusters were generated using hierarchical clustering based on the *d*_*γ*_ distance-matrix. A dendrogram corresponding to the clustering was obtained and used for visualization and annotation of clusters with iTOL^42^.

### G. Modeling hATG8–pSAR complexes and identification of LIR–LDS interactions

We assembled a list of hATG8–pSAR pairs with experimental evidence of physical interactions and modeled their 3D complex structure using AlphaFold2-Multimer^43^. We obtained several models corresponding to each complex using default parameters for the number of templates, energy tolerance, and structure relaxation parameters. The database used to obtain homologous sequences and template structures was last updated on 01/01/2022. We analyzed the 25 top-ranked models by building contact maps. Experimentally resolved structures of hATG8–SAR+ complexes, obtained from RCSB PDB, were mapped to UniProt sequences to compute structural coverage.

Residue-wise contacts were computed with in-house scripts using MDAnalysis^44^ *AB*_cnts_ = Σ_*i*∈*A*_ Σ_*j*∈*B*_ σ (|*r*_*ij*_|), where the sums extend over heavy-atom positions of the interacting residues (*i, j*) and *σ*(|*r*_*ij*_ |) = [1 *−*0.5(1 + tanh(|*r*_*ij*_|*−*6))], which is a smooth sigmoidal counting function to limit interactions below cut-off distance *r*_*ij*_ *≤*6 Å. Contact maps were averaged over all 25 AlphaFold models to obtain interacting sites on pSARs and hATG8 proteins.

### H. Identification of cargo components of pSARs

We computed cell-line expression similarity 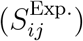 using Spearman’s rank correlation coefficient (normalized to [0, 1]) of expression (in transcripts per million) of proteins (*i, j*) in 1055 cell-lines obtained from the Human Protein Atlas^31^. We computed subcellular localization similarity 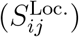 using cosine similarities of binary vectors describing subcellular localization. We used weights 1.0 and 0.3 to denote primary and secondary locations and 0.0 for the absence of proteins (*i, j*) in specific subcellular compartments as obtained from the Human Protein Atlas^31^. Network proximities of proteins (*i, j*), 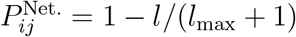, were evaluated after computing the number of intermediate nodes (*l*) in the shortest path connecting them within the physical PPI networks obtained from combining BioGRID^36^ and STRING^35^. Networks were visualized and rendered using Cytoscape^45^.

## III. RESULTS

### A. Autophagosomal protein-inventory reveals purged processes and pathways

A total of 4627 proteins were detected in the content profiling of autophagosomes. We found a substantial overlap (*∼*96%) in the identity of proteins detected within intact autophagosomes isolated under basal autophagy condition, upon EBSS-mediated starvation, and MG132-induced proteasome inhibition (Fig. 1a), indicating that a substantial protein portfolio of autophagic vesicles is preserved and robust under different conditions. However, the relative abundance of these proteins could be variable.To identify autophagosomal protein contents specific to EBSS and MG132 treatment, we compared their autophagosomal protein-profiles to basal conditions (Fig. 1b). Consistent with previous findings^23^, we found that EBSS treatment did not alter the protein-profile significantly. Compared to basal conditions, 16 proteins involved in cytoplasmic and mitochondrial ribosome biogenesis showed an enhanced degradation rate, indicating that energy and resource-rich processes are selectively purged under starvation (Fig. S1a). However, proteasome inhibition with MG132 treatment resulted in increased localization of 147 proteins. These proteins predominantly correspond to the 20S proteasomal complex, protein inclusions found in neurodegenerative disease pathways, and viral infection pathways (Fig. S1b).

**FIG. 1.**
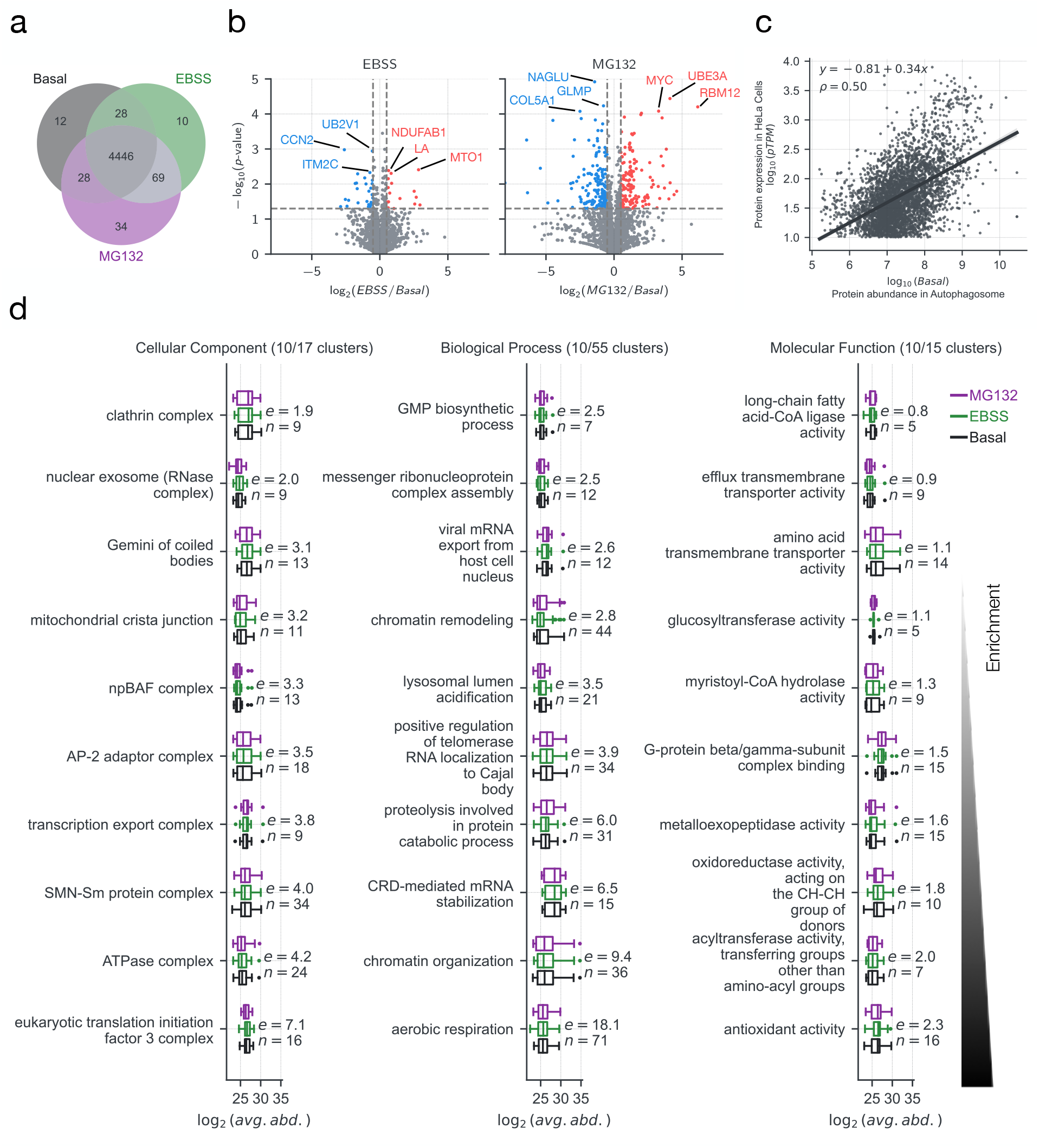
Autophagosomal protein inventory reveals purged processes and pathways. (**a**) Venn diagram showing a 96% overlap in the identity of proteins detected using content-profiling of autophagosomes extracted from HeLa cells under non-treated (Basal; grey), starved (EBSS; green), and proteasome inhibited (MG132; purple) states. (**b**) Volcano plots showing up-regulated (red) and down-regulated (blue) proteins under treatment with EBBS and MG132. (**c**) A scatter plot showing the correlation between HeLa cell protein expression (pTPMs, protein transcripts per million) and relative protein abundance in purified autophagosomes (basal conditions). (**d**) Clustering of 3000 most abundant proteins in autophagosomes at the three gene ontologies (CC, BP, and MF). The top 10 enriched functional clusters (vs. human proteome) and their relative abundances (boxplots) in autophagosomes are shown with the cluster size, n, and enrichment score, e, respectively.

To test if the protein turnover in autophagosomes is controlled at the transcriptional level, we compared the protein expression levels (protein transcripts per million, pTPM) detected in HeLa cells with their relative protein-abundances measured in vesicles isolated from HeLa cells (Fig. 1c). We found a positive correlation (Pearson *ρ* = 0.5), indicating that there was a proportional representation of proteins within autophagosomes under basal conditions, demonstrating that there were no biases in the proteomics data of intact autophagic vesicles. To identify recycled components, purged cellular processes, and molecular functions, with high turnover, we performed GO functional clustering of the 3000 most abundant autophagosomal proteins (Fig. 1d). We found that the autophagosomes are primarily populated by large multi-subunit protein components corresponding to proteins with essential housekeeping functions., i.e., gene transcription, protein translation, splicing, mitochondrial respiration, and critical components of the endocytic pathway. To identify whether the treatment conditions affected functional clustering, we also clustered the 500 most abundant proteins detected in autophagosomes extracted under all three conditions independently. Although we did not find any clear detectable differences in the GO functional clusters of proteins during basal and starvation conditions (Fig. S2a vs. S2b), we found that MG132-treatment enriched proteins corresponding to proteasomal core-components and proteins involved in proteasome-mediated protein catabolism (Fig. S2c), indicating a major rerouting of protein catabolism using autophagy, confirming previous analysis^23^.

### B. Identification of new potential SARs

The presence of an LC3 interacting region, (LIR) or GABARAP interacting motif (GIM) is the hallmark of SARs. We scanned the sequences of all proteins detected in autophagosomes (PTMX) for conserved LIR/GIM-like sites to identify potential new selective autophagy receptors (Fig. 2a). We detected a total of 44 847 LIR sites within the 5032 proteins identified within autophagosomes. We then compared the distribution of confidence scores (PSSM log-odds ratio) of these LIR segments with the distribution of scores for LIR sites detected in the gold standard positive (GSP) datasets. On average, the confidence scores of LIRs for GSP (SAR+) and GSP (SAR–) proteins were higher than that of the PTMX proteins. Using a PSSM cut-off (red line, Fig. 2a; PSSM score *≥* 16 (median value for LIRs of GSP (SAR+)), we identified 1892 high-scoring LIR sites corresponding to 1401 potential selective autophagy receptors (pSARs).

**FIG. 2.**
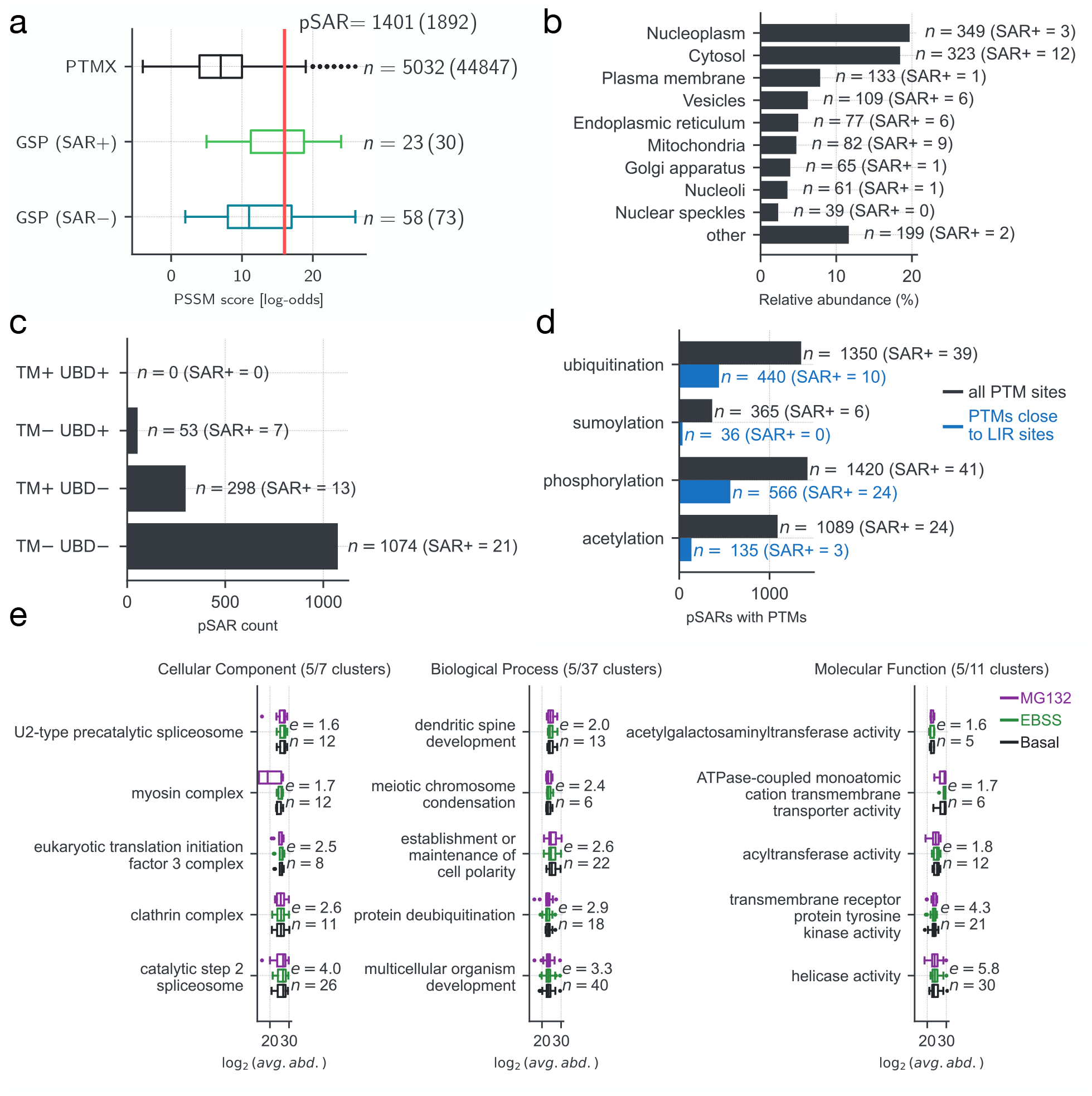
Identification of potential SARs. (**a**) The distribution of PSSM scores for LIRs was obtained by scanning PTMX and GSP datasets using iLIR. Proteins with high-confidence LIR predictions (*n* = 1401, PSSM *≥* 16; red line) and SAR+ (41) proteins present in the PTMX dataset are grouped as pSARs (1425). (**b**) The primary sub-cellular location of pSARs, weighted by their relative abundance in autophagosomes, shows the diversity of selective pathways operating under basal conditions. (**c**) Categorization of 1425 pSARs by likely mode of action: solution phase/ membrane-bound and involvement in ubiquitin-dependent/ independent pathways. (**d**) PTMs on pSARs, close to predicted LIR sites (*±*5 residues), influence their function and regulate selective autophagic pathways. (**e**) Top 5 clusters from GO functional clustering analysis of pSARs.

To identify the cellular compartments that are predominantly targeted by pSARs, we mapped their primary subcellular location along with their relative abundance in autophagosomes (Fig. 2b). We found that the cytosol, nucleoplasm, and endocytic membranes cumulatively account for *∼*73% of the observed abundance of pSARs within autophagosomes. Further, we found that multiple well-known SAR+ proteins (*n* = 17) were also picked up in our expanded pSAR list (Fig. 2b).

To determine if these pSARs target solution-phase or membrane-bound cargo components, we annotated them with consensus transmembrane predictions. To determine if they participated in ubiquitin (Ub)-dependent or Ub-independent pathways, we additionally mapped the various ubiquitin-binding domains (UBDs) within these proteins (Fig. 2c). We found *∼*75% and *∼*20% of pSARs target solution-phase cargo and membrane-bound cargo components, respectively, using Ub-independent pathways. Only a meager 3% of pSARs showed explicit UBDs and were potentially able to direct solution-phase components to autophagosomes.

Next, to test if the pSAR functions could also be regulated by possible PTMs that are frequent to selective pathways we mapped known experimentally detected PTM sites onto pSAR and SAR+ sequences (Fig. 2d; *n* = 1425). We found phosphorylation, followed by ubiquitination and acetylation, are the three dominant PTM in pSARs and SAR+ proteins. We also found multiple PTMs either on the predicted LIR sites or within its close proximity (*±*5 residues) on pSARs (440, 36, 566, and 135 unique pSARs with ubiquitinations, sumoylations, phosphorylations, and acetylations, respectively), indicating that the local density and sequence of PTMs could determine local structure and provide an additional regulatory layer on pSAR functions.We re-analyzed biological processes and molecular functions selectively targeted by pSARs (Fig. 2e). We detected 37 enriched clusters spanning diverse processes essential to HeLa cell physiology. We also found several proteins annotated primarily as transmembrane receptors, kinases, transporters, ion channels, and nuclear proteins, indicating that they could also moonlight and function as pSARs under specific physiological conditions.

### C. Relationship of pSARs and SAR+ proteins

To determine whether the pSARs could directly bind to hATG8-family members, we mapped and filtered physical binding partners of pSARs and SAR+ proteins from global protein–protein interaction networks with experimental evidence (Fig. 3a). We found that 159/1401 pSARs showed evidence for physical binding to at least one of the six hATG8 proteins. We also mapped 27/41 hATG8–SAR+ interactions, indicating that the current PPI databases covered a substantial set of previously well-characterized interactions (65%). Further, upon closer examination, we were able to identify 17 well-known SAR+ proteins in the pSAR list with high-scoring LIR sites. 11 well-known SAR+ proteins were also detected in the larger PTMX dataset, albeit with lower confidence (PSSM score *<* 16). Additionally, we analyzed previously identified SAR+ proteins, their LIR motifs, and available evidence to mediate hATG8 interactions, along with their representation in our PTMX dataset (Tab. S1).

**FIG. 3.**
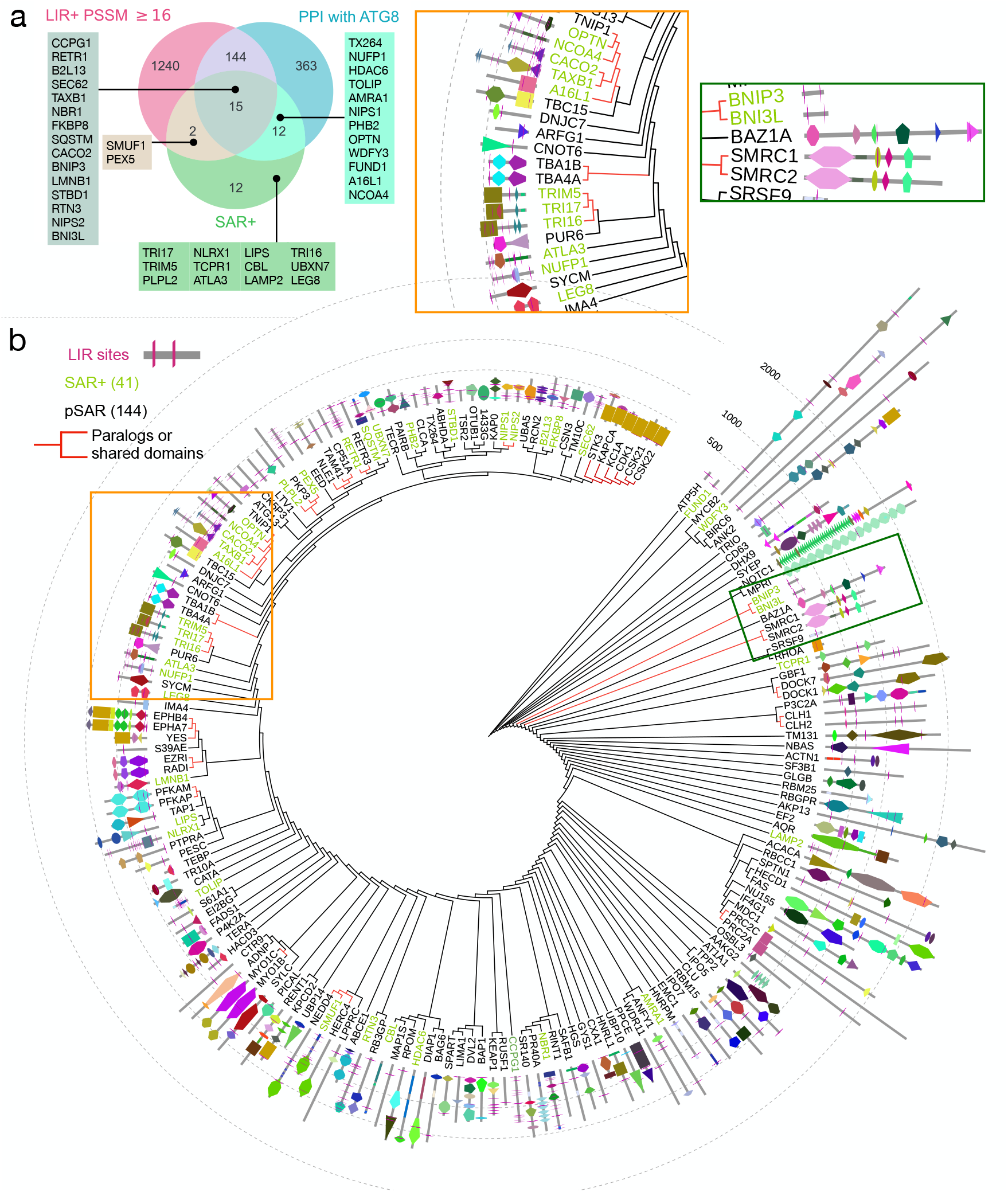
Relationship of pSARs and SAR+ proteins. (**a**) A Venn diagram showing the overlap of autophagosomal proteins (PTMX) with high-confidence LIR-predictions (LIR+ PSSM *≥* 16, pink circle), PTMX proteins interacting with hATG8 (blue circle) and well-established SAR+ (listed proteins in green circle). (**b**) Dendrogram showing the results of hierarchal clustering of pSARs (*n* = 144, black labels) and SAR+ proteins (*n* = 41, green labels) based on distances estimated from pairwise-global alignments (see Methods). The leaves show the mapping of domain architecture from InterPro (filled shapes) and LIR predictions from iLIR (purple) onto each sequence. Concentric circles show protein length around the dendrogram. Insets (orange and green boxes) show a zoom-up of the dendrogram with examples (red branches) of paralogs or proteins with shared domains and sequence elements.

To study the relationship between the newly identified pSARs (144) and the well-known SAR+ (41) proteins, we performed pairwise global alignments to estimate relative distances between proteins and used hierarchical clustering to draw new relations (Fig. 3b). We mapped the domain architecture information and high-scoring LIR-positions onto each pSARs and SAR+ sequence and found that most proteins are dissimilar, showed diverse domain architectures, and had potential LIR sites scattered along their sequences to varying degrees. This indicated a diversity in the landscape of pSARs employed in selective autophagy pathways. Interestingly, we also found several nodes rooting more related proteins (leaves of red branches), i.e., paralogs or proteins with shared domains/sequence elements (orange and green insets in Fig. 3b). Often, one paralog was already annotated as SAR+ (green labels), and the other as pSAR (black), corroborating functional similarity (see Fig. S3 for expanded view). For example, we identified HERC4 and NEDD4, co-clustered with SAR+ protein SMURF1, known to function in xenophagy^46^. In another case, we identified RETR3 (FAM134C), a dominant paralog of the well-known selective ER-phagy receptor RETR1 (FAM134B)^21^, which was only recently demonstrated to function as an independent SAR^47^. These observations demonstrate that mining and effective integration of PPI networks can result in high-quality pSAR predictions, despite datasets not being updated with recent experimental evidence.

### D. pSAR–hATG8 interactions

To validate the interactions of pSARs with hATG8 proteins obtained from the orthogonal mapping of PPI networks and to recognize their role in selective autophagy, we modeled the 3D structures of hATG8–pSAR complexes using the AlphaFold2-Multimer^43^. We were able to successfully model 53 out of *∼*160 complexes (Tab. S2). We made an initial assessment of our iLIR predictions by analyzing 3D structures of hATG8–SAR+ binary protein complexes from the PDB. We found several high-scoring LIR segments were engaged in canonical LIR– LDS binding mode as represented by two examples, NBR1 and CALCOCO2 LIRs (Fig. 4a). Motivated by these findings, we performed a similar analysis with pSARs and analyzed the bound conformations from their modeled complex structures. We found characteristic LIR– LDS binding mode analogous to SAR+ proteins validating our iLIR predictions for pSARs. Amongst the 53 hATG8–pSAR complexes modeled, 27 showed explicit LIR–LDS binding modes, validating 18/103 high-scoring LIR predictions (Tab. S2). We highlight two representative examples, LTV1 and PAIRB, that show the conserved binding mode along with residue-wise contact maps highlighting the LIR–LDS interaction (Fig. 4b). Human ATG8 proteins share similarities with ubiquitin structure, and, in principle, can bind to ubiquitin interaction motifs (UIMs) of pSARs using their UIM-docking site (UDS), a hydrophobic pocket on the opposite face^9^.Although we did not predict UIMs in pSARs and SAR+ proteins, we analyzed modeled complexes for explicit binding at UDS of hATG8 proteins. We found that 37/53 modeled complexes displayed close UIM–UDS contacts (Fig. 4c), indicating an expanded interaction potential of pSARs. In fact, 17/53 modeled complexes display dual interactions, LIR– LDS and UIM–UDS in their bound states. For example, top-ranked models corresponding to LTV1–LC3A complex show both LIR–LDS and UIM–UDS interactions simultaneously. Analysis of all the intermolecular contacts showed that most complexes shared several interfacial residues, indicative of other possible alternate binding modes (Tab. S2).

**FIG. 4.**
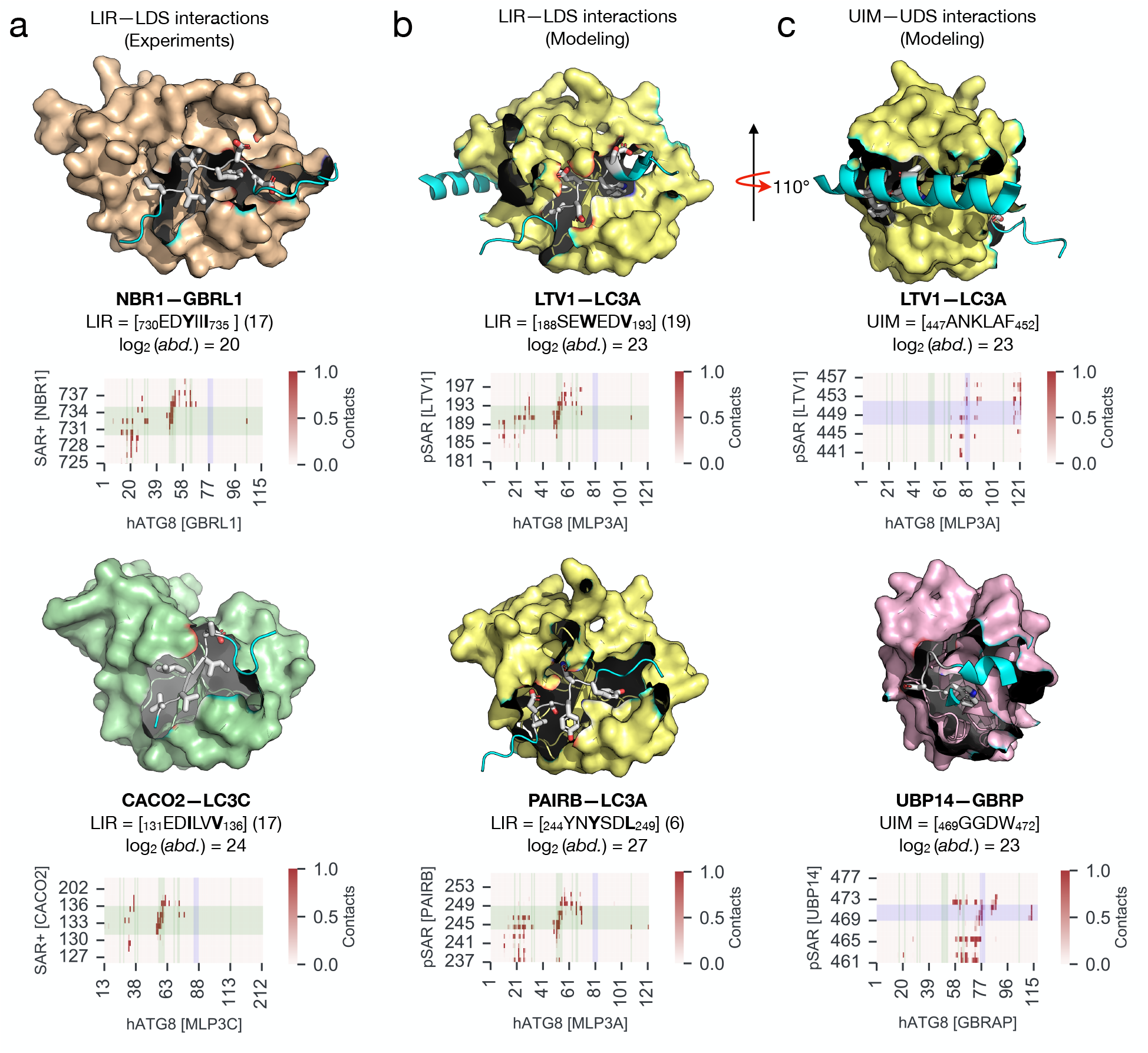
Structure of pSAR–hATG8 complexes. Analysis of 3D structures of hATG8–SAR binary complexes obtained from the PDB and de novo modeling using AlphaFold2-Multimer reveals several examples of the characteristic LIR-LDS and the UIM–UDS binding modes of SARs. (**a**) LIR–LDS interactions of well-known SAR+ proteins, NBR1 and CALCOCO2 from experimentally resolved structures (PDB codes: 2L8J and 3VVW) match our LIR predictions. (**b**). LIR–LDS interactions of new pSAR proteins, LTV1, and PAIRB from our top-ranked AF models. (**c**) Additional UIM–UDS interactions are also observed in AF models of the ATG8-complexes of LTV1 and UBP14. These binding modes are consistent with our LIR predictions (sequence with PSSM scores in parentheses; side chains shown as sticks in 3D models) for SAR+ and pSAR proteins (cyan cartoon) and plug the hydrophobic pockets (HP1 and HP2) on hATG8 (shown as colored surface). Contact maps averaged over 25 models of each binary complex highlight the consistency of LIR– LDS (green-shaded) and UIM–UDS (blue-shaded) interactions in multiple modeling iterations.

### E. Cargo components targeted by pSARs

We detected 5273 proteins in the purified autophagosomes (PTMX; see methods). These potential cargo proteins are targeted by either selective or non-selective pathways into autophagosomes. To determine whether some proteins were selective autophagic cargos and were detected in intact autophagosome-proteomics in large quantities (PTMX) as a result of direct recruitment by pSARs, we estimated the interaction potential of pSARs. We chose the 500 most abundant cargo proteins (non-pSARs and non-SAR+ proteins) and quantified their likelihood to interact with pSARs and SAR+ proteins. Based on the assumption that co-expressed and co-localized proteins are more likely to form direct physical connections and assemble in close proximity, typical of autophagic cargos, we computed cell-line co-expression similarity, subcellular localization similarity, and PPI network proximity of pSARs and SAR+ with potential cargo proteins. We estimated 92 500 similarity and proximity measures corresponding to every SAR–cargo pair (Fig. 5a, 185 *×*500 pairs). The heatmap displays relationships for only the 50 most abundant cargo proteins (basal condition) with 25 SAR+, and 25 pSARs, respectively.

**FIG. 5.**
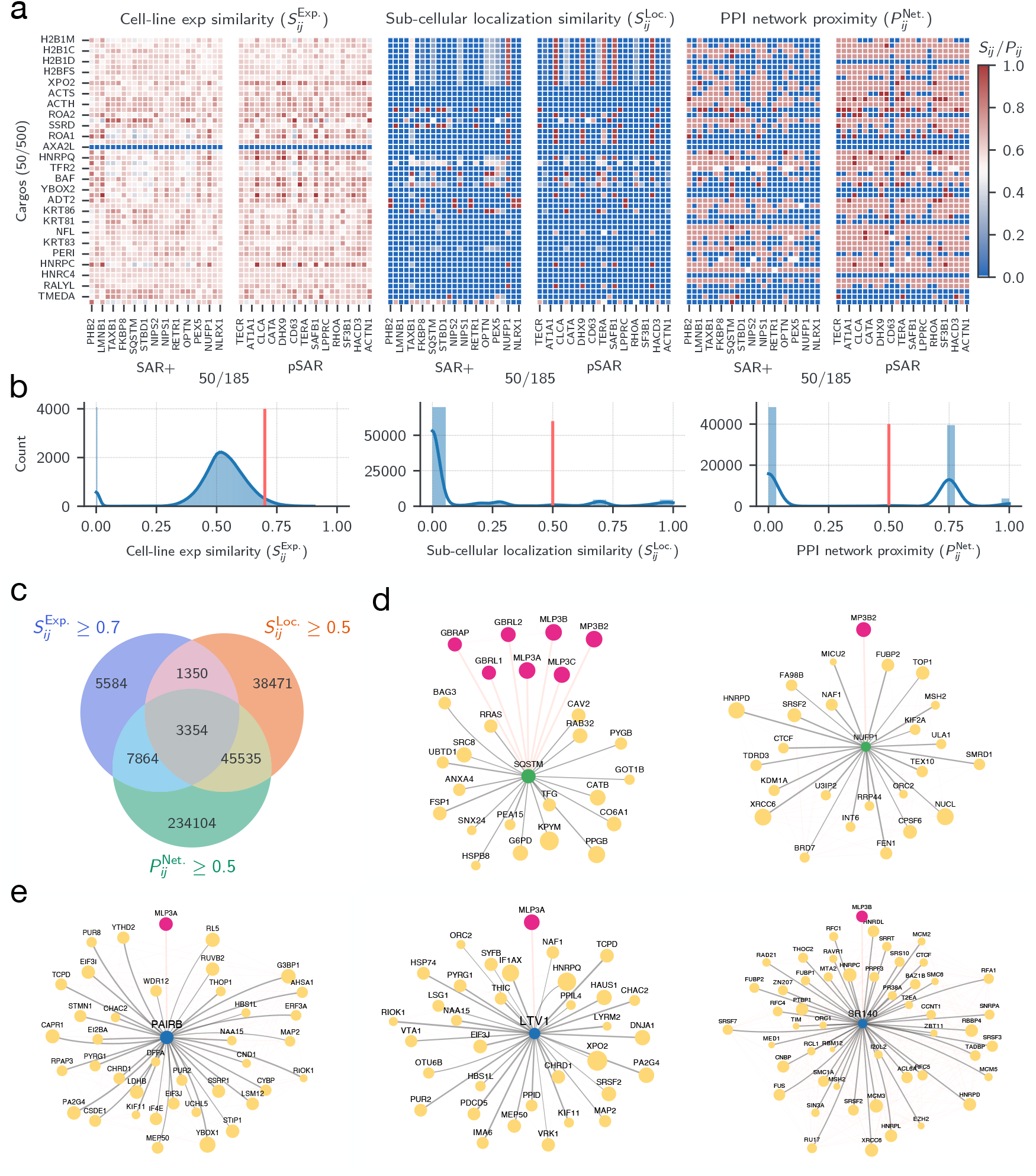
SAR–cargo interactions. (**a**) Representative heat maps showing the co-expression, colocalization, and network proximity of SARs and cargo proteins found in PTMX. The map shows the 50 most abundant cargo proteins, along with 25 SAR+ and 25 pSAR proteins. (**b**) Histograms show the distributions of the similarity and proximity measures within the entire dataset (185 *×* 500 pairs) with threshold values (red line). (**c**) Venn diagram showing the filter process used to obtain high-confidence pSAR–cargo pairs. Physical interaction networks of (**d**) two SAR+ proteins (green), SQSTM and NUFP1, and (**e**) three new pSAR proteins (blue), PAIRB, LTV1, and SR140, reveal high-confidence specific cargo molecules (yellow) targeted for degradation. Network nodes are scaled by protein abundances detected in autophagosomes, and edges are weighted by the sum of similarity and proximity measures computed for each pSAR–cargo pair. Background physical interactions between proteins (red edges), and interactions of pSAR with hATG8 proteins (purple), are shown (red edges with increased weight).

We found that the cell-line co-expression similarities for these corresponding pairs were continuously distributed and relatively high, whereas the distribution of the subcellular location similarity index and PPI network proximity index was more discrete with few peaked distinct values (Fig. 5b). By filtering protein pairs using threshold scores and by combining the three measures, we added additional confidence to the likelihood of direct interactions. We found 3354 unique, high-confidence interactions between SAR–cargo pairs and detected 15 SAR+ proteins and 80 pSARs with at least 5 or more cargo molecules in the autophagic vesicles (see Fig. 5c and Tab. S3). Mapping this information onto the entire physical PPI network revealed the direct sub-set of cargo molecules, which are co-expressed, co-localized, and in close proximity with each SAR+ (Fig. 5d; 2/15 SAR+, SQSTM, NUFP1 shown) and pSAR proteins (Fig. 5e; 3/80 new pSAR, PAIRB, LTV1, and SR140).

## IV. DISCUSSION

Various cellular homeostatic pathways converge, and their perturbation results in the induction of autophagy. Although the degradative pathway is active under basal (normal) conditions, stress conditions such as nutrient deprivation, hypoxia, oxidative stress, and infections induce autophagic activity, increasing its flux and result in a substantially enhanced cargo turnover^48^. Autophagosomes act as sinks, collecting cellular material and designated cargos for degradation by employing bulk and selective autophagy pathways. The extent and diversity of non-selective and selective pathways operating under any given cellular state are large, and often their relative contributions towards enhanced autophagic activity under stress are difficult to decouple.

The FACS-based isolation and proteomics profiling of autophagosomes^23^ offers a unique opportunity for “dumpster diving” into cellular garbage bins. This novel isolation is an *ad hoc* method, without genetic manipulation, resulting in high quantities of pure and intact autophagosomes, preserving the autophagic history. By isolating and profiling a large ensemble of intact autophagic vesicles from cells—approximately 8 million autophagosomes analyzed using proteomics for each given replicate and treatment condition—we obtain space- and time-averaged sampling of autophagosomal contents. This provides a good vantage point for identifying previously unknown proteins associated with autophagy and enables the filling of knowledge gaps.

By directly looking into autophagosome contents to identify new SARs, we constrained our search space, removing a large number of false positives often found in screens for novel hATG8 binders. We predicted *∼*1400 potentially new SARs containing high-scoring LIR sites. By effectively combining LIR predictions and direct hATG8 binding, we report *∼*185 proteins from our dataset that could function as SARs. These proteins display all essential characteristics of SARs. The identification of several known SARs amongst them adds tremendous value, increasing the confidence in our predictions. Further, the mapping of these proteins to diverse subcellular compartments, the establishment of known physical interactions with hATG8 proteins, and their direct detection in autophagosomes in large abundance strengthen their functional role as SARs. Furthermore, we identified several molecular features of these proteins: preservation of LIR sites among paralogs with known SAR functions, positioning of TM regions and UBDs to detect the likely mode of action (solution phase-/membrane-mediated and Ub-dependent/-independent mechanisms), and their PTMs (required for SAR regulation), providing substantial evidence for their functioning as SARs.The interaction landscape of hATG8 proteins and SARs is vast. Motif-mediated binding is the hallmark of these low-affinity interactions. By modeling binary complex structures of hATG8 and pSAR using the state-of-the-art AI-based AlphaFold-Multimer, we demonstrated the importance of LIR–LDS binding modes consistent with iLIR predictions. We also found additional UIM–UDS binding modes in our modeled complexes that were not explicitly predicted. It is important to note that we identified hATG8 family members engaged in interactions with LIR-containing proteins in non-autophagic processes, i.e., SAR– proteins with LIR motifs. Furthermore, we found several examples of low-confidence LIR sites (PSSM *<* 16), directly engaged in interactions with the LDS. These are examples of atypical or non-canonical LIR sites, and information on many such sites with critical functions is still emerging^9^. These findings suggest that key motif-based interactions essential for SAR binding (LIR/LDS and UIM/UDS) are already captured by the AlphaFold neural network. In fact, a recent study used protein modeling using Alphafold-Multimer to identify both canonical and atypical AIM/LIR motifs with a high level of accuracy^49^. Therefore, our models provide direct mechanistic examples for testing the recruitment of the phagophore and autophagic machinery by these pSARs.

Despite identifying pSARs, their cargo specificities, and recruitment mechanisms are often untested, even for well-established selective autophagy pathways. By integrating orthogonal measures on protein co-expression, co-localization, and network proximity, we identified proteins closely associated with the new pSARs. These molecules could be associated with autophagosomes due to direct recruitment by SARs, or indirectly. Given that they are detected in large abundances directly within autophagic vesicles along with pSARs, they are likely to be recruited cargo components. The number of SARs functioning in any given cell is vast. A diverse cargo portfolio demands selective signaling and recruiting mechanisms to fine-tune the autophagic response. Further, the landscape of active SARs and cargo components changes drastically in various cell types and tissues, indicating tight regulatory control mechanisms^50^. By mining the autophagosomal protein inventory, we have evaluated the autophagic state of HeLa cells in response to treatments and identified several important selective autophagy factors. Our analysis provides several lines of thought for hypothesis generation, along with clear examples for testing by experimentation. It remains to be seen how these data can be leveraged to enhance our understanding of selective autophagy pathways.

## V. ACKNOWLEDGMENTS

We thank Christian Behl and Ivan Dikic for their advice and technical support, Anne-Claire Jacomin for critical reading of the manuscript, and David Krause for system administration. This work is supported by funds from the CRC project on Selective Autophagy (Project-ID 259130777-SFB1177). A.C., A.D., and S.A.P-C. are partially funded by EU-bOPEN, PROXIDRUGs, and EnABLE consortia, respectively. We also thank the Center for Supercomputing, Goethe University Frankfurt for computing time on the Goethe-HLR cluster.

## VII. AUTHOR CONTRIBUTIONS

A.C. analyzed the proteomics datasets and performed all the computational predictions and analyses with assistance from A.D., S.A.P-C., and support from R.M.B. A.C. and R.M.B. conceived and designed the research, performed the analysis with help from A.K., and wrote the paper with input from all the authors.

## VIII. DATA AVAILABILITY STATEMENT

The data that support the finding of this study are available from the corresponding author upon reasonable request.

## X. SUPPORTING INFORMATION

Additional supporting information file containing Figures S1 and S2 is available as a separate pdf. Additional data files corresponding to Tables S1–S3 are available as xlsx files.

