## Supplemental Figs 1-2 for "Identification of potential selective autophagy receptors from protein-content profiling of autophagosomes"

### Supplementary Figures and Tables

This pdf contains Supplementary Figures (Figs. S1 and S2)

All Supplementary Tables are provided as separate xlsx files.

**Table S1** (Table_S1_extended_pSAR_info.xlsx): List of pSAR and SAR+ proteins with LIR predictions.

**Table S2** (Table_S2_AF_modeling_summary.xlsx): Summary of AlphaFold modeling and analysis of pSAR–hATG8 complexes.

**Table S3** (Table_S3_pSAR_cargo_info.xlsx): List of pSAR and associated cargo proteins.

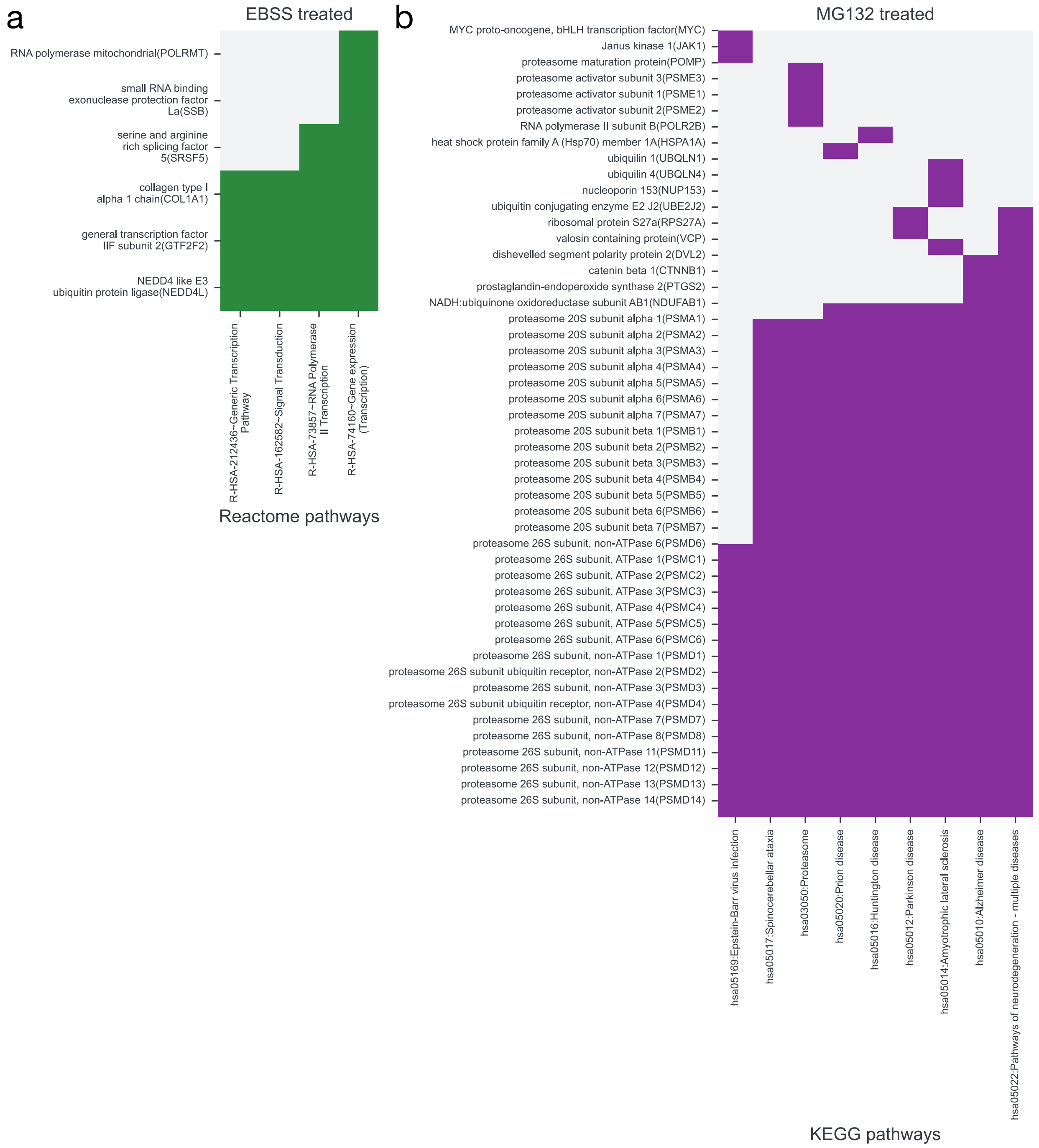


FIG. S1. **Effect of starvation and proteasome inhibition.** Pathway enrichment analysis of up-regulated proteins found in autophagosomes after (a) EBSS treatment (6/12 enriched proteins linked to 4 gene expression pathways) and (b) MG132 treatment (49/147 enriched proteins associated with 9 infection/disease pathways linked to protein aggregates).

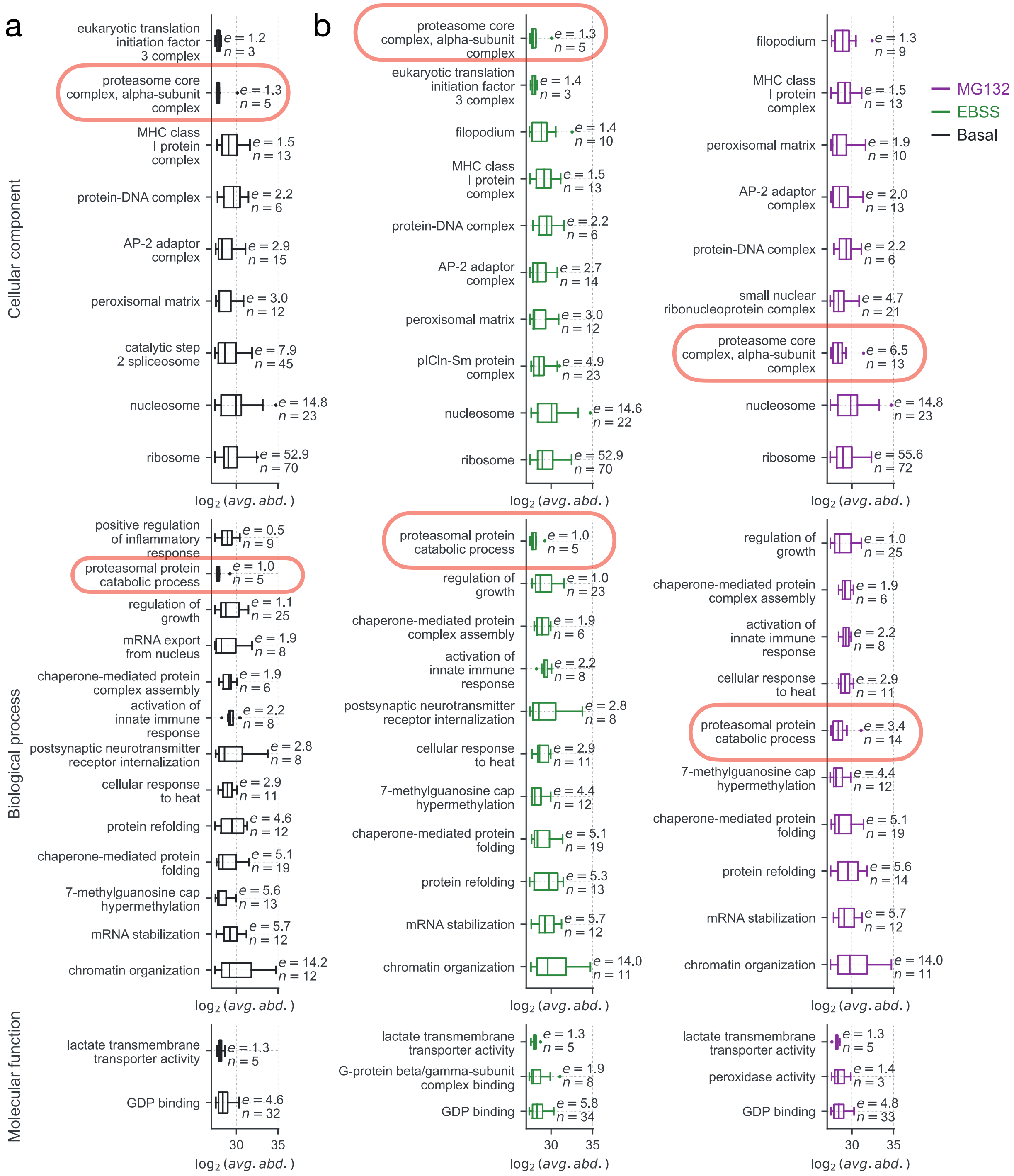


FIG. S2. **Treatment-specific changes in protein-inventory of autophagosomes.** Functional clustering of 500 most abundant proteins detected in autophagosomes extracted from HeLa cells under (a) basal, (b) EBSS, and (c) MG132 treatment. Enriched functional clusters (vs. human proteome) obtained at the three gene ontologies (CC, BP, and MF) are shown along with their relative abundances (box plots), cluster size, n, and enrichment score, e, respectively. Note that MG132 treatment enriched proteins forming the core-proteasomal complex and proteins involved in proteosome-mediated catabolic processes.
